# Echoes from the last Green Sahara: whole genome analysis of Fulani, a key population to unveil the genetic evolutionary history of Africa

**DOI:** 10.1101/2023.04.06.535569

**Authors:** Eugenia D’Atanasio, Flavia Risi, Francesco Ravasini, Francesco Montinaro, Mogge Hajiesmaeil, Biancamaria Bonucci, Letizia Pistacchia, Daniel Amoako-Sakyi, Maria Bonito, Sara Onidi, Giulia Colombo, Ornella Semino, Giovanni Destro Bisol, Paolo Anagnostou, Mait Metspalu, Kristiina Tambets, Beniamino Trombetta, Fulvio Cruciani

## Abstract

**Background:** The Sahelian Fulani are the largest nomadic pastoral ethnic group. Their origins are still largely unknown and their Eurasian genetic component is usually explained by recent admixture events with northern African groups. However, it has also been proposed that Fulani may be the descendants of ancient groups settled in the Sahara during its last Green phase (12000-5000 BP), as also suggested by Y chromosome results.

**Results:** We produced 23 high-coverage (30 ×) whole genomes from Fulani individuals from 8 Sahelian countries, plus 17 samples from other African groups and 3 Europeans as controls, for a total of 43 new whole genome sequences. These data have been compared with 814 published modern whole genomes and analyzed together with relevant published ancient individuals (for a total of > 1800 samples). These analyses showed that the non-sub-Saharan genetic ancestry component of Fulani cannot be only explained by recent admixture events, but it could be shaped at least in part by older events by events more ancient than previously reported, possibly tracing its origin to the last Green Sahara.

**Conclusions:** According to our results, Fulani may be the descendants of Saharan cattle herders settled in that area during the last Green Sahara. The exact ancestry composition of such ghost Saharan population(s) cannot be completely unveiled from modern genomes only, but the joint analysis with the available African ancient samples suggested a similarity between ancient Saharans and Late Neolithic Moroccans.

## Background

About one third of the African continent is occupied by the Sahara desert that spans from the Atlantic coast to the Red Sea and represents the widest hot desert on Earth. At the north, the Sahara is bordered by the Mediterranean Sea, while on the south the Sahelian belt separates the desert from tropical forests further south. Because of its geographic position and ecological features, the Sahel is characterized by an intermediate environment mainly composed of savannah and grasslands. The northern part of the Sahelian belt shows a semi-arid climate, where no extensive agriculture is possible, while the more humid southern area allows agricultural practices. However, pastoralist groups can exploit the pastures in the northern Sahel and then move southwards during the driest months. So, the Sahel represents a natural contact area between the pastoralist and agricultural lifestyle, represented by groups speaking languages from three out of four African linguistic families, namely Afro-Asiatic, Nilo-Saharan and Niger-Congo [1].

Among the people living in the Sahel, the Fulani, speaking a language belonging to the Niger-Congo linguistic family, represent an interesting case. They are the largest pastoral ethnic group in the world, with an estimated population of 20-40 million people settled in a broad area covering 18 African countries, from the Atlantic coast to the lake Chad basin and further east to the Blue Nile region in Sudan [2]. Fulani are historically nomadic pastoralists, although most of them have now adopted a sedentary lifestyle based on farming. Nowadays, most Fulani live in western Africa, where the first evidence of their presence traces back to the XI century, when they settled on the Fouta Djallon highlands in central Guinea. Later, they moved eastward along the Niger river and they arrived at the lake Chad region in the XV century [3,4]. Despite their recent history being relatively known, their origins are still a matter of debate. Over time, it has been proposed that their ancestors could be ancient Egyptians, Nubians, Persians, Jews, Arabs, Ethiopians or western Africans; however, the most widely accepted theories trace the Fulani origins back to ancient northern Africans or near eastern populations [5–8]. In this context, it has also been proposed that the Fulani may be the descendants of ancient Saharan populations [1]. Indeed, the Sahara has not always been as harsh as today: between 12,000 and 5,000 before present (BP), it was a lush and fertile environment occupied by savannah, forests and a wide hydrogeographic network of rivers and lakes. This phase, generally known as “Green Sahara”, was just the last of several alternating arid and humid phases that have characterized the geological history of this region [9,10]. This region was inhabited by different human groups with peculiar material cultures during the last Green Sahara period, as suggested by several paleoanthropological and archeological findings [9], including several Saharan rock paintings as the rock art in the Tassili-n-Ajjer plateau in the Algerian Sahara, dated to 8,000 BP. Interestingly, this painting represents cattle and pastoralism rituals very similar to the ceremonies still practiced by present-day Fulani [1,11], suggesting a possible link between these people and the ancient inhabitants of the Sahara.

The ancient Saharan hypothesis about the Fulani origins seems also to be supported by some analyses of the human Y chromosome: Fulani show a high proportion of a Y lineage, i.e. E1b-M2/Z15939, dating back to the last Green Sahara. This lineage is observed at lower frequencies also in western and northern Africa, while it is absent in other sub-Saharan regions, pointing to a Green Saharan origin in the Fulani ancestors [12]. However, other genetic data seem to give contrasting results. Considering the other human uniparental system, i.e. the mitochondrial DNA (mtDNA), Fulani harbor a non negligible proportion of Eurasian U5b and H1cb haplotypes, possibly arrived from a northern African source [13,14]. As for the autosomal DNA, genome-wide studies have shown that Fulani people display both a western African component and a northern African/Eurasian one, but the estimates of such components differ across different studies and sub-groups [15–19]. In this context, the most striking result has been obtained from the lactase locus: Fulani show a high proportion of the lactase persistence allele T-13910 [20], that is typical of the European populations [21–23]. This finding has been explained suggesting that the European lactase persistence haplotype was first introduced in northern Africa and then in the Fulani population by recent admixture with a northern African population about 2000 years ago [15]. Similarly, the presence of European ancestry components on the autosomal or uniparental portions of the Fulani genome is usually explained by recent admixture between a Western or Central African source and an “Eurasian” source dating back to the last two millennia [18].

Despite the general agreement about the presence of a “non-western-African” component in the Fulani genome, the origin and the extent of such a component is still a matter of debate. In the last decades, the increasing number of published individuals analyzed by whole genome sequencing (WGS) has allowed to shed light on different dynamics of the populations included in such studies [24–28]. However, the number of whole genomes from the African continent is disproportionately low compared to the sequences from other continents, with only few African individuals included in the main genome variation projects [29–31], despite the importance of Africa for the past and recent human history. Recently, some projects have focused on the African whole genome variability, partially filling this gap [19,32–36]; however, only few whole genomes from Fulani individuals have been published and analyzed so far [19,34,37]. The results from these researches confirmed the presence of a “non-western-African” component, that has been suggested to be acquired in recent times, possibly from an eastern Afro-Asiatic source [19,34,38]; however, the contrast with data from other portions of the genome still remains and the questions about the origin and past history of Fulani people are still open. To address these issues, we analyzed 23 high-coverage (30 ×) whole genomes of Fulani from 8 countries, that have been compared with additional 17 samples from other African ethnic groups, plus 3 European samples as controls, for a total of 43 whole genome sequences presented here for the first time. These data have been then compared with whole genomes available from the literature [19,30,31,39] from relevant ethnic groups and geographic areas. The modern dataset has been then analyzed together with relevant published ancient individuals [40] to obtain a detailed description and accurate time estimates of the main population dynamics during the Fulani history and about the peopling of the African continent and the role of the last Green Sahara in general.

## Results

### Whole genome diversity and population structure of the Fulani people

We selected 23 Fulani samples from 8 countries in the Sahelian belt. To this group of samples, 17 samples from other African populations and 3 European individuals have been added, for a total of 43 samples, representing 14 different populations, 9 African linguistic sub-families and 13 sampling countries, analyzed by high coverage (> 30 ×) whole genome sequencing (Table 1). The 43 samples presented here were merged with 771 published whole genome sequences (including 2 additional Fulani from Cameroon) from four different projects, namely the study by Fan and colleagues [19], the Simons Genome Diversity Project (SGDP) [30], the high-coverage sequences from the 1000 Genomes Project [39] and the whole genomes from the Human Genome Diversity Projects (HGDP) [31]. The published samples have been selected to represent relevant ethnic groups and geographic regions for our study (Additional File 1, Supplementary Table S1) to create a modern dataset including a total of 4,143,934 markers (1,907,678 after LD pruning) from 814 individuals (Fig. 1).

**Table 1:**
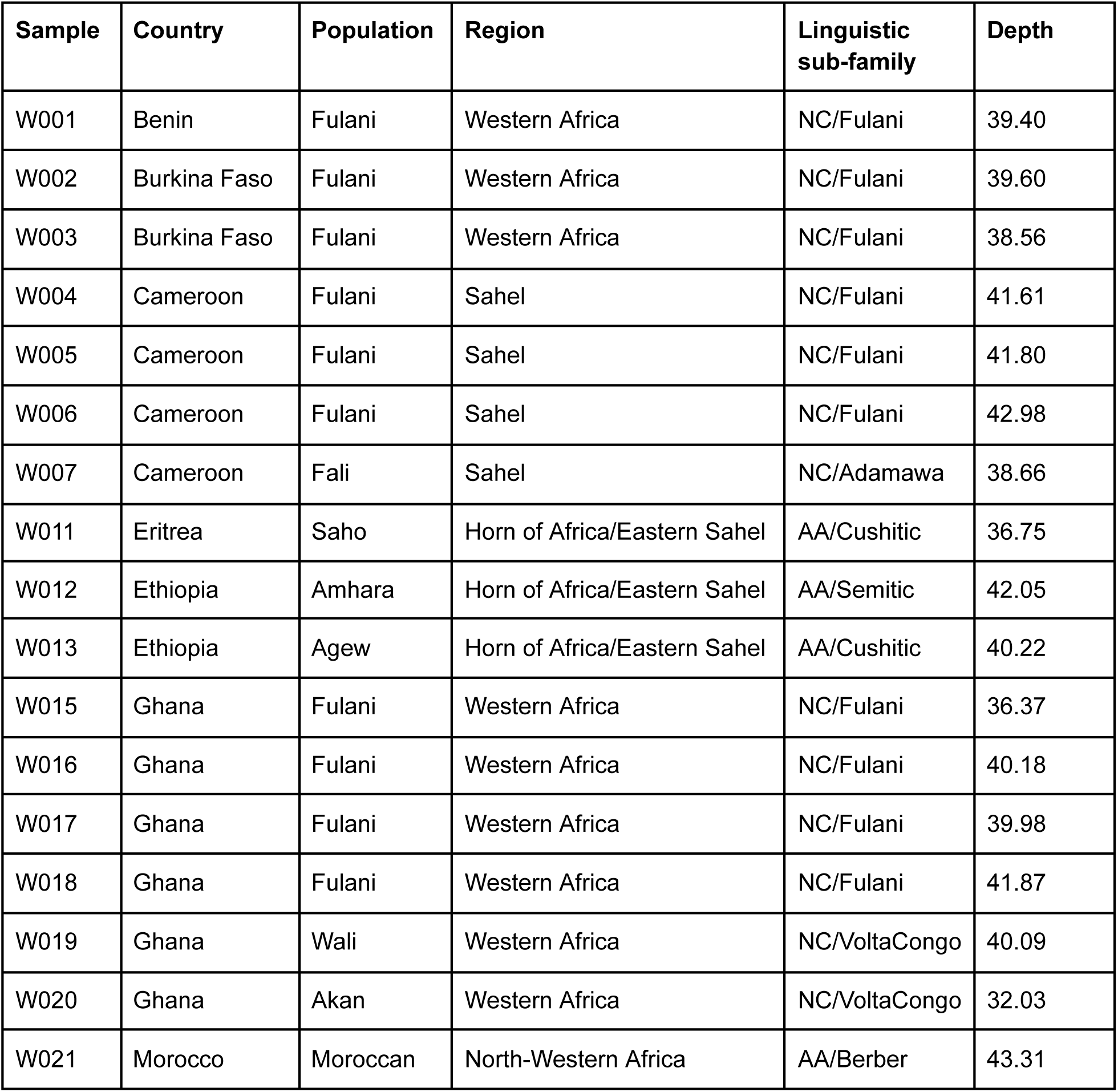
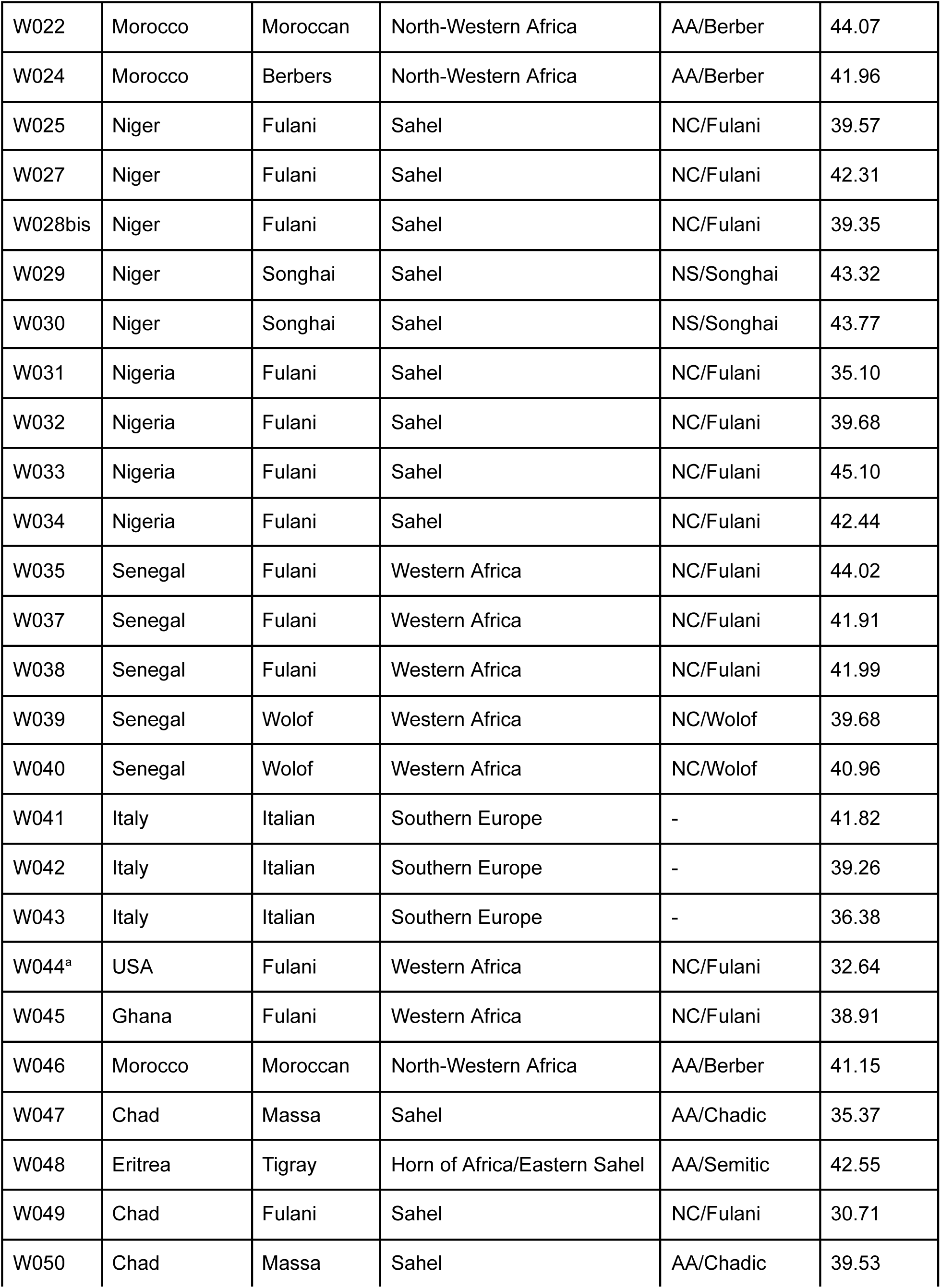
Description of the individuals sequenced in this study. For each sample, the country, the population, the region, the linguistic affiliation and the average sequencing depth is reported. The language affiliation is reported for African groups only. a. For this Fulani individual, the region was assigned based on his declared place of origin in Guinea. AA: Afro-Asiatic; NC: Niger-Congo; NS: Nilo-Saharan (more information is reported in Additional File 1, Supplementary Table S1).

**Figure 1:**
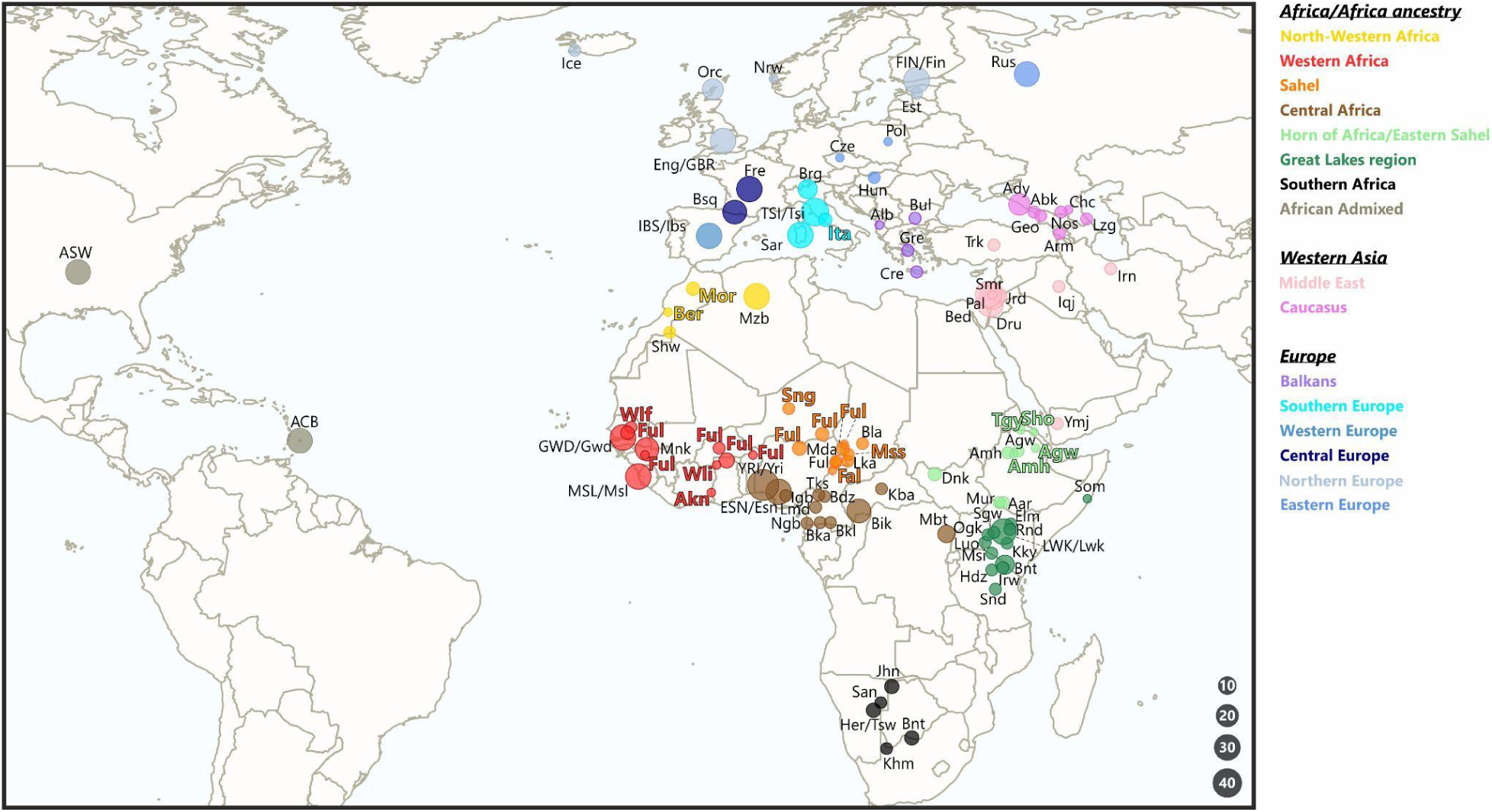
Map of the sampling locations of the individuals analyzed in this study. Each pie represents a population, is drawn proportionally to the sample size (legend at the bottom right) and is coloured according to the geographical region (legend at the top right). Population acronyms are described in Additional File 1, Supplementary *Table S1*. Groups sequenced in this study for the first time have coloured labels according to the geographical region. CEU from 1000 Genomes have not been represented on this map.

**Figure 2:**
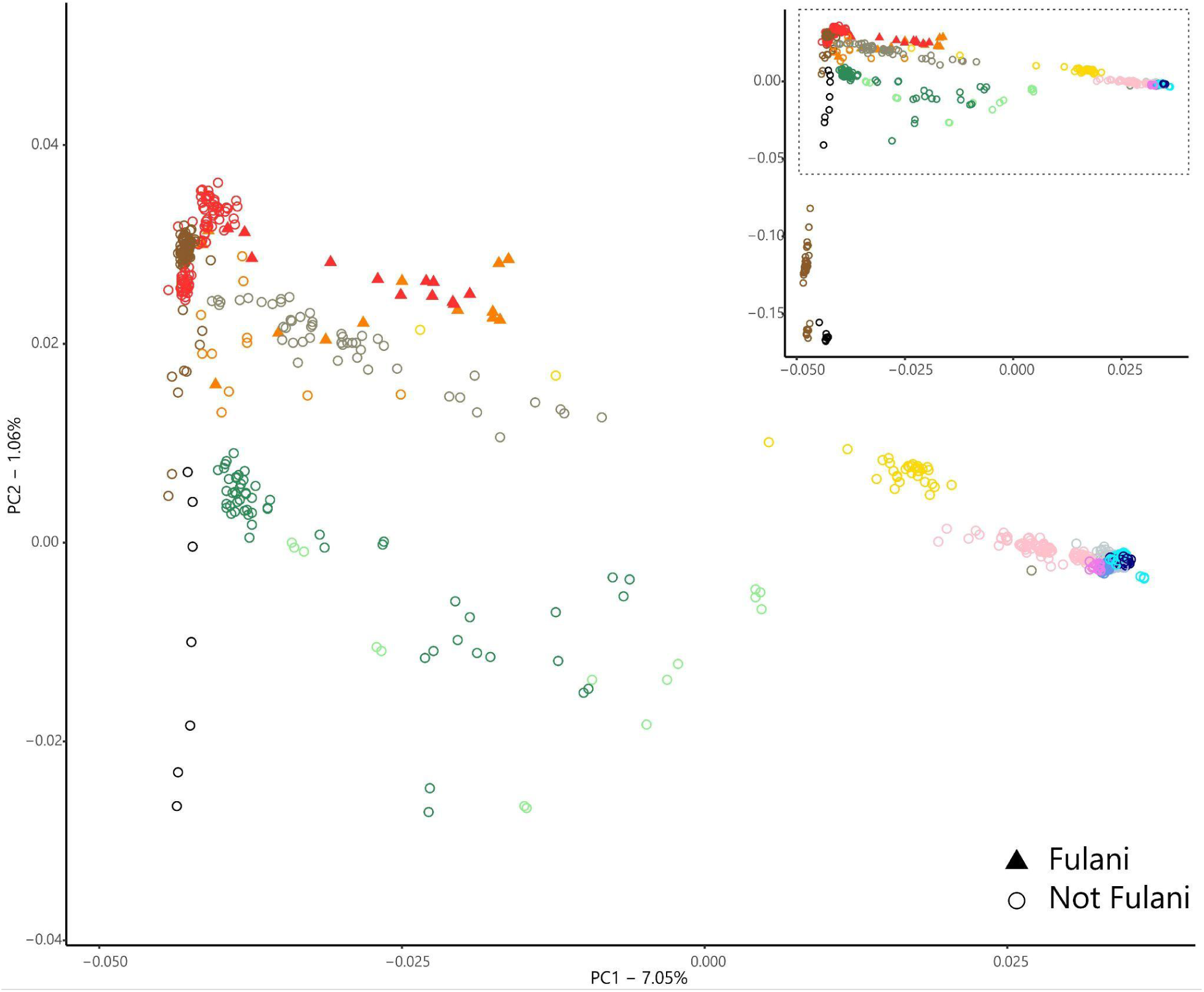
Principal component analysis (PCA) of the modern dataset. Each symbol represents an individual, with shapes assigned to distinguish the Fulani individuals (legend in the bottom right) and coloured according to the geographical region as in Fig. 1. The main plot represents a magnification of a subset (dashed rectangle) of the total PCA in the inset. An interactive version of the total PCA with sample-by-sample labels is available as Additional File 2.

First, we performed a principal component analysis (PCA, Fig. 2): in general, most of the variability can be explained by geographic location and ethnic group affiliations, with the first PC separating the southern African Khoe-San speaking individuals and some central African Rainforest hunter-gatherers from all the others, while the second PC separates Africans and non Africans, according to previous studies [18,30].

Looking at the PCA in more detail, we can observe that three groups of samples form long stretches from central/western African cluster to northern African and Eurasian clusters: 1) the eastern African samples (shades of green); 2) the African admixed (grey); 3) the Fulani individuals (red/orange triangles).

The eastern African group is the most dispersed along both PC1 and PC2, with people from the Great Lakes region (dark green) nearer to the Bantu groups (including Luhya) and people from the Horn (light green) shifted towards the Middle East/Europe along PC2, consistently with their known Eurasiatic ancestry component [41]. The second group is composed by the two African admixed populations, African Caribbean in Barbados (ACB) and African in Southwest US (ASW) from the 1000 Genomes project [29]: consistently with their known history related to the trans-Atlantic slave trade, these samples occupy a long stretch along PC2, with some individuals tightly clustering with western/central African people and some others shifted towards the Europeans, while their variation is very low along PC1 [42]. Finally, the third group is represented by the Fulani people (orange/red triangles). Differently from the two previous groups, the Fulani cluster is more homogeneous along both PCs, occupying a defined region in the PCA space, between western/central Africa and the non-African regions along PC2. Along the PC1, the position of the Fulani cluster is above the line of the northern African sample, differently from eastern African and African admixed groups that lie below. This observed difference between the Fulani and the other two groups with a well-known Eurasian component may suggest a different source of the non-sub-Saharan ancestry in the Fulani genomes, in addition to differences between the groups contributing to their sub-Saharan ancestry. Despite the Fulani general homogeneity, it is worth noting that not all the individuals fall in the Fulani cluster: indeed, some of them fall in the western/central African groups or among the African-admixed samples, suggesting that admixture events between Fulani and different neighboring populations may have shaped their genetic pool at some extent [15,43].

We further investigate the population structure of the Fulani people compared to the other African and non-African groups by performing an admixture analysis [44], for K from 2 to 10 (Additional File 3, Supplementary Figure S1). For the sake of clarity, hereafter we refer to the genetic ancestry components with the name of the regions/populations mainly representing them. At K=2, the African (blue) and non-African (orange) ancestry components are clearly separated, while K=3 distinguishes the African hunter-gatherer ancestry (yellow), as previously observed [45]. At K=4, when the cross validation error minimizes, we can see a northern African/Middle Eastern ancestry component (light blue), different from the European one: interestingly, this component is observed both in Fulani and in the eastern African groups, that have a known admixture history with the northern Africa/Middle East [46,41]. On the other hand, this ancestry component is absent in the African Admixed ACB and ASW [29], which instead show a non-African ancestry of European origin. At K=5, we can observe an eastern African ancestry component (dark blue), different from the northern African/Middle Eastern one and present in the groups from eastern Africa. At K=6, the northern African ancestry component (purple) is distinguished from the Middle Eastern one: among the Fulani, this new component replaces the old one and represents the second most frequent component (37% on average). Higher Ks further define the Fulani sub-Saharan ancestry component (violet), that ca be defined as a Senegambian component since it is representative (∼ 100%) of these people (i.e. Wolof, Mende and Mandenka), while it decreases south-and eastward. This admixture analysis further confirms the presence of two subgroups among our Fulani samples, clearly differentiated by their ancestry composition. On the basis of both the PCA and admixture results, we then subset the Fulani in two groups: hereafter, FulaniA and FulaniB will refer respectively to the samples within the main Fulani PCA/admixture cluster or outside it (Additional File 1, Supplementary Table S1).

### New clues on Fulani origins from ancient DNA

In order to shed light on the origin of Fulani, especially of their non-sub-Saharan ancestry component, we merged the modern dataset described above with the Allen Ancient DNA Resource [40], selecting only the ancient and modern samples from the same macroregions as the individuals in our modern dataset (Additional File 1, Supplementary Table S2). First, we performed a PCA by projecting the ancient samples on the principal components obtained only from the modern samples (Additional File 4). The PCA recapitulates what has been observed from Fig. 1, with ancient individuals mainly falling into the genetic variability ranges of modern samples from the same regions.

We then performed an admixture analysis on the same dataset (Additional File 3, Supplementary Figure S2). Focusing on the Fulani groups, we can see that the FulaniA cluster is differentiated from other sub-Saharan groups and from the FulaniB since K=2, showing an average 25% of the non-sub-Saharan ancestry component (azure), similarly to the African Admixed groups. This non-sub-Saharan component is further defined with increasing K values; at K=5 (lowest cross validation error), we can see that the FulaniA cluster shows 3 non-sub-Saharan ancestry components: 1) an azure ancestry component, possibly corresponding to the Anatolian Neolithic (AN); 2) a blue ancestry component, representing the Iran Neolithic/Caucasus hunter-gatherers (CHG); 3) a negligible red ancestry component (less than 2%), corresponding to the Western hunter-gatherers (WHG) (Fig. 3).

**Figure 3.**
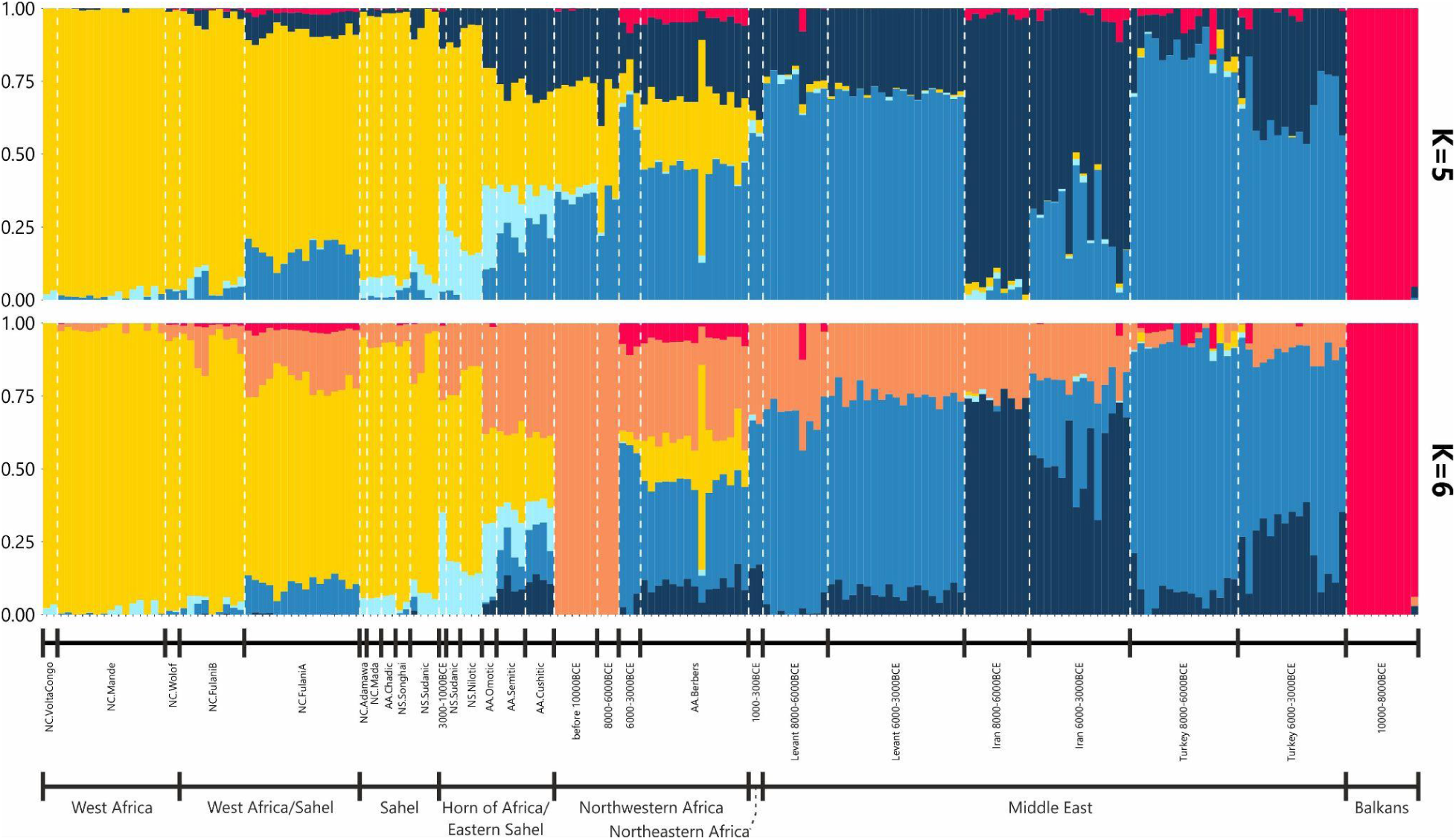
**Admixture analysis on ancient and modern dataset for K=5 and K=6.** Each bar represents a different individual, each color a different genetic ancestry component. Only a subset of individuals has been represented, selecting only relevant groups and randomly subsampling groups larger than 25 (see Additional File 3, Supplementary Figure S2 for the full version of the admixture plot).

The same components are also observed in the FulaniB and in both the African Admixed groups, but at drastically different proportions, suggesting that these groups experienced different evolutionary trajectories. At K=6 (second lowest cross validation error), we observe that the non-sub-Saharan ancestry component in FulaniA is mainly characterized by the new orange component. Interestingly, this component is typical of north-western African individuals older than 8000 BP (more specifically, Taforalt and IAM individuals) [47,48] and can be also observed in other Sahelian/Eastern African groups, while it is virtually absent from African Admixed and other sub-Saharan populations (Fig. 3). On the other hand, the Iranian Neolithic blue ancestry component is not represented anymore among FulaniA. Higher Ks confirmed this pattern where FulaniA share a non-negligible portion of their non sub-Saharan ancestry with ancient northern African people. As for the FulaniA sub-Saharan ancestry component (yellow), this analysis confirmed the link with the Wolof and Mende groups already highlighted by the admixture based on modern data only.

We further explored the link between Fulani (A and B) and ancient people by performing D-statistics in the form D(Fulani, sub-Saharan Africa, Eurasia/North Africa, Chimpanzee), where a) Fulani were alternatively represented by FulaniA or FulaniB; b) excluding the Southern Africa and Northern European regions from sub-Saharan Africa and Eurasia/North Africa respectively; c) testing both FulaniA and FulaniB also as sub-Saharan groups and d) using Chimpanzee as outgroup. Overall, the Eurasia/North Africa pool tended to share more alleles with both Fulani clusters rather than central/western African people and less compared to eastern African populations (both ancient and modern). However, when focusing on the relationships between the two Fulani clusters and the ancient north-western African individuals in the form D(FulaniA, FulaniB, ancient North-western Africa, Chimpanzee), we observed that FulaniA were closer to all the 3 ancient north-western African groups than FulaniB (Additional File 1, Table S3 and Additional File 3, Figure S3). When analyzed by qpWave/qpAdm framework with two putative sources, both Fulani groups were modeled by an ancient sample from Cameroon (about 7000 BP) and the Early Neolithic Moroccans (about 6000 BP), with Iberomaurusian-like component being higher among FulaniA (Additional File 1, Table S3A). In addition, FulaniA were also successfully modeled by modern Berbers and modern Sahelian people. When 3 putative sources were tested, we observed that FulaniA can be only modeled by the same two previous ancient sources plus the eastern Sahelian Agew at approximately the same proportions, while FulaniB could be successfully modeled by two sub-Saharan sources and a northern African one, with the latter accounting for no more than 27% (Additional File 1, Table S3B). Finally, we also performed 10 admixture graphs varying the number of admixture edges from 2 to 5 and retained the three graphs characterized by the lowest score, testing only the Fulani A. For all the inferred graphs FulaniA were modeled as composed of two main ancestries. The first ancestry (between 60% and 70%) is close to Western African groups related to Mende from Sierra Leone (MSL). This group also harbored a minor proportion from a ghost population which stems as an outgroup of all the other populations analyzed, as already suggested in previous studies [49]. The second ancestry (more than 30%) was derived from an unsampled group close to Iberomaurusian samples, modeled as a sister group of ancient Northern Africans, or alternatively, as an outgroup of all the “Eurasian-ancestry” enriched groups (Additional File 3, Supplementary Figure S4). This may suggest that, at least part of the non-sub-Saharan ancestry in present day Fulani might have an ancient origin, and is not exclusively derived by migrations in the last two millennia.

### Patterns of shared haplotypes and runs of homozygosity

To infer patterns of shared ancestry between Fulani groups and other populations, we identified and analyzed the identity-by-descent (IBD) segments longer than 2 cM in our modern dataset (Additional File1, Table S5). FulaniA shared more IBD segments with themselves, while FulaniB shared more IBD with FulaniA and themselves. Interestingly, we could observe a general pattern differentiating FulaniA and FulaniB, with the former sharing more segments with Sahelian, eastern African, northern African and Eurasian populations, while the latter shared more IBD with western-central African groups. When comparing short (< 5 cM) and long (> 5 cM) IBD segments, we could observe that this pattern was mainly driven by the shortest IBD segments, suggesting that it formed in more ancient time, while differences between the two Fulani clusters are more nuanced for the long IBD segments. We further explored the links between Fulani and neighboring sub-Saharan people by performing D-statistics in the form D(Sahel, West Africa, Fulani, Chimpanzee), where Fulani could be alternatively FulaniA or FulaniB (Additional file 1, Supplementary Table S6). Overall, both Fulani clusters were characterized by the same general pattern, although FulaniA showed a slight trend toward Sahelian groups speaking Nilo-Saharan languages, with the notable exception of the Wolof from Western Africa, in line with IBD results.

We then investigated the patterns of runs of homozygosity (ROH) to shed light on the demographic history of the Fulani (Additional File 1, Supplementary Table S7). The two Fulani clusters showed differences, with FulaniB being more similar to other western and central African groups both in terms of number of ROHs and their total length (Fig. 4).

**Figure 4:**
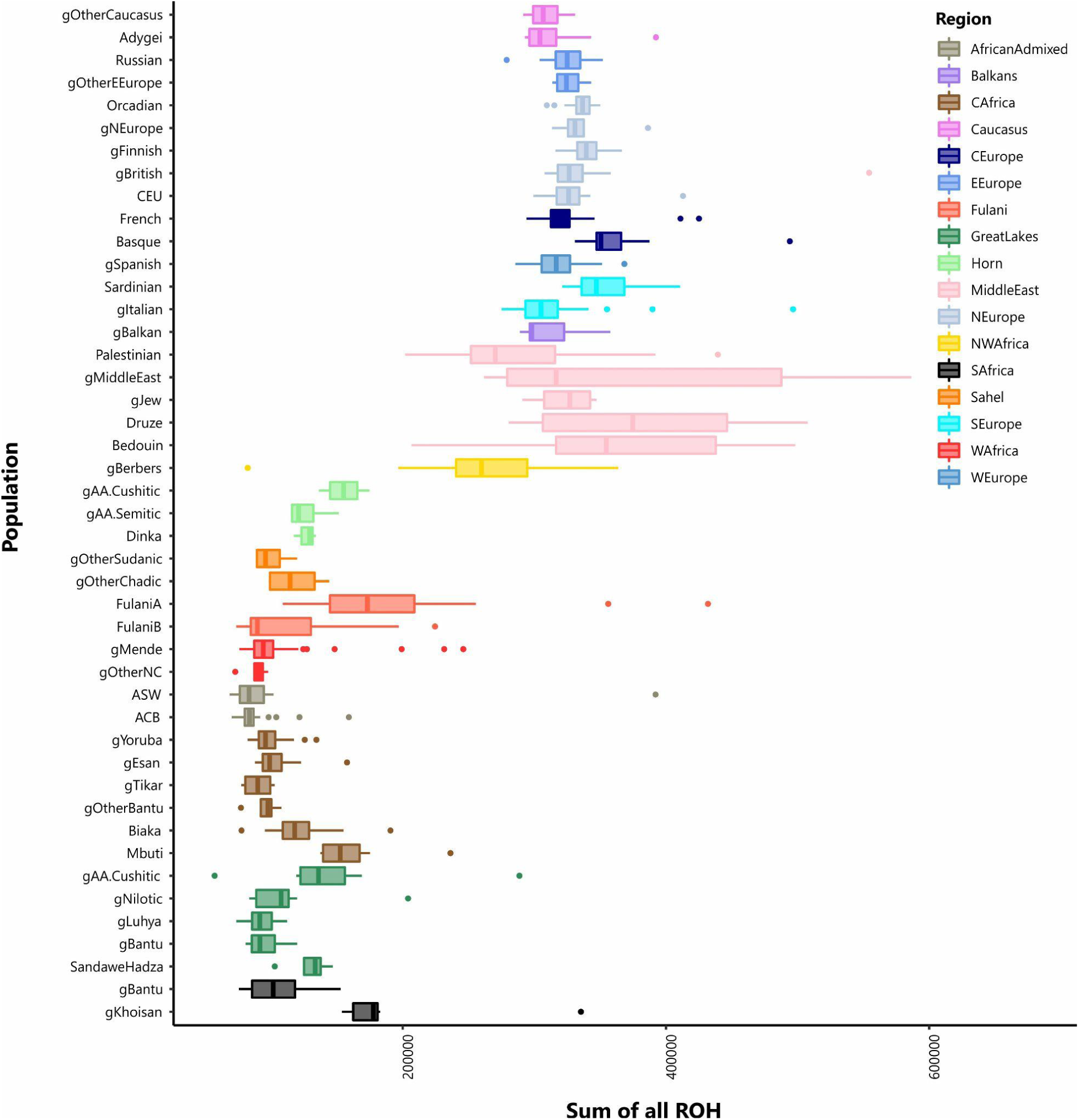
**Analysis of runs of homozygosity (ROH).** Each boxplot summarizes the total length of ROH segments by population group (see Additional File 1, Supplementary Table S1). Observations have been defined as outlier if they were 1.5 times the interquartile range less than the first quartile or greater than the third quartile. Population groups with a sample size less than 3 have not been plotted.

On the other hand, FulaniA showed more ROHs compared to their neighboring groups, with a pattern between Sahelian/central African populations on one hand and northern African populations on the other. We further explored the ROH pattern subsetting the Fulani clusters in Sahelian or western African groups and we observed that FulaniA and FulaniB individuals from western Africa showed very similar ROH patterns (Additional File 3, Supplementary Figure S5A). On the contrary, Sahelian FulaniA showed a higher number of ROHs; when analyzing the short (< 1.5 Mb) and long (> 1.5 Mb) ROHs separately (Additional File 3, Supplementary Figure S5B-C), we observed that this pattern was mainly driven by the long segments. On the other hand, the short ROH segments recreated the clear differentiation between FulaniA and FulaniB, regardless of their sampling region, with the former showing the number and the mean sum of ROHs more similar to eastern Africans rather than western/central African groups.

## Discussion

### A Green Sahara story

Most recent studies based on the genome-wide variability suggested that the origin of the non-sub-Saharan ancestry of Fulani traces back to relatively recent times (1800-300 BP), after admixture events between a western African and northern African/southern European source [15,18,37,38]. Here, we analyzed the Fulani genomes in the frame of the worldwide genetic variability considering both modern and ancient individuals. In this context, the analysis of aDNA allowed us to characterize the non-sub-Saharan ancestry component in the Fulani in more detail, shedding new light on its origin. By comparing the Fulani genetic diversity with those of the large number of ancient individuals currently available [40] through an admixture analysis (K=6, Fig. 3), we observed that their non-sub-Saharan ancestry is characterized by a component observed in extremely high proportions (virtually 100%) also in ancient Morroccans older than 8000 BP [47,50], and an azure Levantine component. We could also see a non-negligible fraction of the WHG red component, as in the Late Neolithic Moroccan individuals, possibly arrived from Iberia to the Maghreb [47]. On the other hand, Fulani lacked the Iran Neolithic/CHG blue component, which was observed instead in modern northern Africans, Middle Easterners and Europeans. This pattern of ancestry components can be hardly explained solely by recent admixture between western African and northern African groups. Indeed, in this case, we would have expected to observe also the Iran Neolithic/CHG blue component in the Fulani, since this component is present in all the modern northern African, European and Middle Eastern samples. So, the admixture event(s) that forged the non-sub-Saharan ancestry in Fulani should have occurred before the arrival of this component in the areas to the north of the Sahara. It is challenging to obtain exact time estimates for such events [51], considering that they were probably followed by later admixture events involving the same or very similar source populations [15,18,37,38]. However, considering the results of the admixture analysis (K=6), we can try to define broad time windows. Indeed, considering the presence of the azure (Levantine) and red (WHG) components in Fulani, in addition to the orange (Iberomaurusian) one, and the absence of the blue (Iran Neolithic/CHG) component, we can postulate that their non-sub-Saharan component dates back to about 8000-7000 years ago from a source population similar (except for the blue component) to the Late Neolithic Moroccans (dated about 5000 BCE, considering their radiocarbon time estimates) (Additional 2, Supplementary Table S2). In addition, the qpAdm analysis also points to a period corresponding to 7000 BP considering the radiocarbon time estimates of the two ancient sources that successfully modelled both FulaniA and FulaniB (Additional 2, Supplementary Table S4). Intriguingly, this time window corresponds to the period of the last Green Sahara.

The Sahara has played and still plays an important role in the population history of the African continent. Currently, it acts as a strong geographic barrier because of its arid environment; however, during its humid phases (called Green Sahara), it became a fertile and lush land inhabited by different species, including humans [10]. The last of these fertile periods occurred approximately between 12000 and 5000 BP, during the Holocene climatic optimum, and the archaeological findings from that time are characterized by different material cultures [9], testifying the rich diversity of the populations in that region. Indeed, the fertile environment of the Sahara, with extensive hydrogeological networks connecting rivers and lakes, may have promoted extensive contacts between the different groups settled in this area. The last Green Sahara was also characterized by social and cultural changes, such as the spreading of pastoralism and farming, both developed locally or arrived from the Middle East to northeastern Africa between 7000 and 6000 years ago [1,52,53].

In this scenario, the formation of the non-sub-Saharan component in Fulani should not be interpreted as a simple admixture event between a western African source group and a northern African one. On the contrary, we should consider that many different groups inhabited the Saharan area at that time [53,54] and the favorable climatic conditions could have allowed a relatively continuous gene flow between them (Fig. 5), as also suggested by the presence of a western African ancestry component in the Iberomaurusians [50] and in addition to the main northeast-to-northwest and northeast-to-east routes [48,55,56].

**Figure 5:**
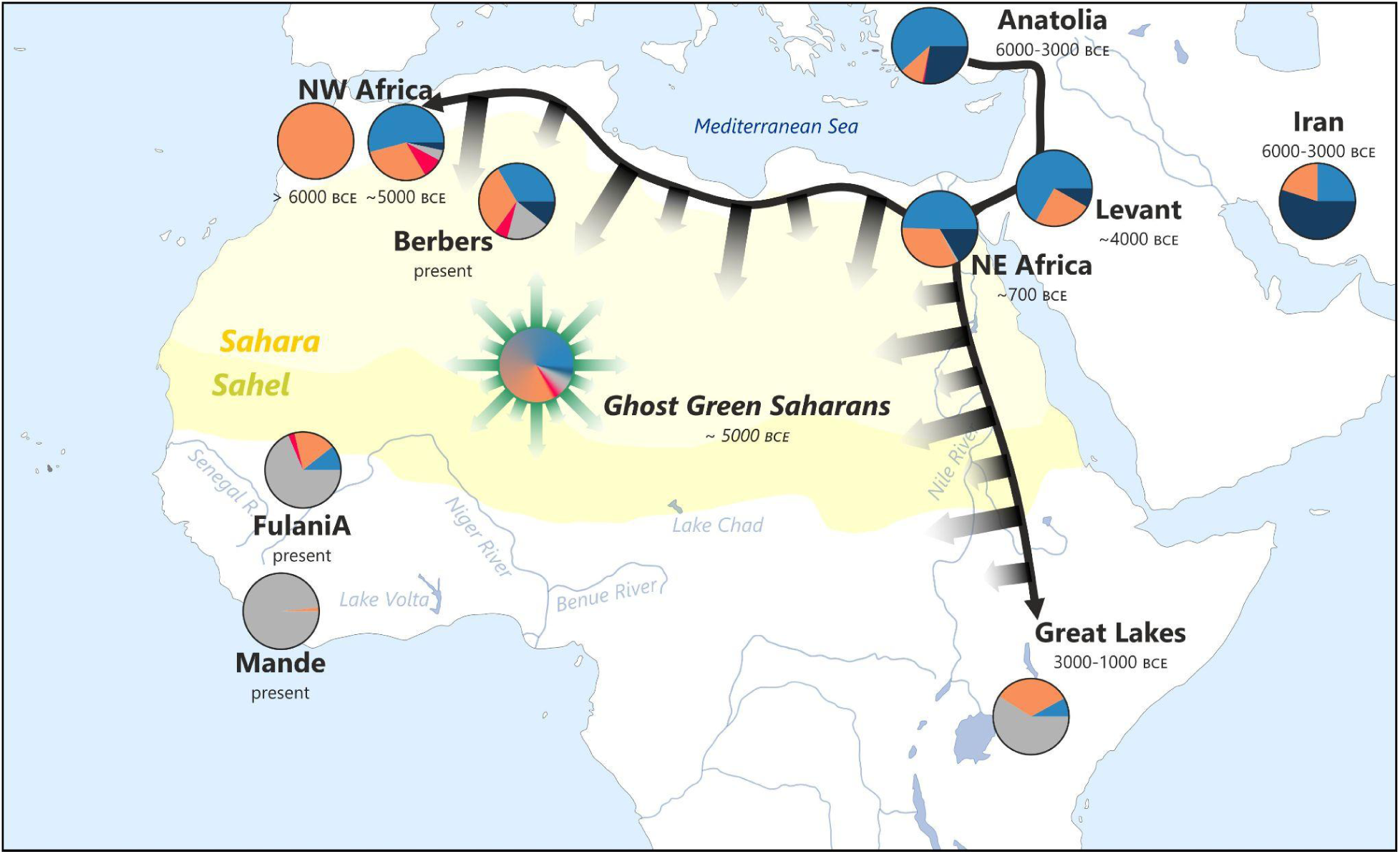
Movements of main non-sub-Saharan ancestry components in the Green Sahara 8000-7000 BP and a possible scenario for the formation of the Green Saharan population(s). The ancestry components are the same represented in the admixture with modern and ancient individuals for the same populations (Fig. 3 and Additional File 3, Supplementary Figure S2). Faded arrows represent possible movements across the Green Sahara.

These Saharan people may have actually been the ancestors of present-day Sahelian people, including Fulani. This is also supported by our admixture analysis, showing that also the Neolithic pastoralists from Kenya and the modern central Sahelian groups showed the orange and azure components at non-negligible proportions (Fig. 3 and Additional File 3, Supplementary Figure S2). Moreover, the absence of the blue component in the modern Nilo-Saharan groups from both central and eastern Sahel further confirms this hypothesis, since it has been proposed that the Nilo-Saharan languages were much more widespread in the Green Sahara region during the last humid phase, to be replaced by Afro-Asiatic languages only in more recent times (about 1500 years ago) [9,53]. In this context, it is also worth noting that a recent whole genome study further confirmed the presence of a shared ancestry between Fulani and Afro-Asiatic speakers from Eastern Africa [37]. Interestingly, it has also been proposed that the Green Sahara area (more precisely the lake Chad basin, occupied by the Megalake Chad at the time [9]) was also the homeland of the Niger-Congo (i.e. the language currently spoken by Fulani) about 7000 years ago, intriguingly corresponding to the possible time estimate of the Fulani non-sub-Saharan components. This language then spread westwards (and eastwards to a lesser extent) replacing the pre-existing languages [57].

The exact genetic ancestry composition of the “ghost” Saharan populations cannot be assessed without ancient individuals from that area at that time; however, on the basis of the ancient data currently available, we can propose that they were genetically similar to the Late Neolithic Moroccans here analyzed, although this group already shows the Iranian Neolithic/CHG component that was probably more diluted further south, while the yellow western African component was possibly present at higher proportions (Fig. 5).

With the end of the Green Sahara and the subsequent gradual desertification of that region, the Saharan groups moved westwards, eastwards, southwards or northwards, as also suggested by the Y chromosome data [12]. This phenomenon was not abrupt and was slightly longer in the central Sahel rather than in the east depending on the local hydrogeological conditions; according to the archaeological evidence, the changing climate conditions led to changes in the socio-economic organization of the different groups. In particular, it has been suggested that cattle pastoralism became the predominant form of subsistence at that time since it was a more reliable source of food, leading to the establishment of a pan-Saharan cattle cult, as testified by rock paintings and rituals that spread westwards with the pastoral groups in search of new pastures [54,58]. This scenario may also reconcile the apparent contrasting evidence from different genetic systems linking Fulani with sub-Saharan, northern African, European or eastern African groups [12,15,17,37,38,43,59–61]; indeed, if we assume the past presence of Green Saharan populations with extensive contacts, it is not surprising that these groups could share genetic affinities. With the Sahara desertification and the following different population dynamics, the genetic differentiation between the different groups may have led to the loss or maintenance of different genetic variants because of genetic drift and/or selection.

All this bulk of data seems to point to a Green Sahara origin of the Fulani with regard to their non-sub-Saharan component. Their ancestors were possibly Saharan cattle herders that moved westwards in response to the changing climate and then mixed with local people. Later, the peculiar Fulani lifestyle, historically characterized by nomadism and endogamy [59,62], and their population size dramatic decrease [37] could have prevented the dilution of this ancient “Green Saharan” ancestry component that can be still identified.

Finally, both Fulani clusters show the same ancestry components, pointing to a common ancestry of the two groups, although at different proportions, suggesting different population dynamics.

### Demographic history of Fulani

All the analyses performed in this study highlighted the presence of two distinct Fulani clusters, here defined FulaniA and FulaniB. The sub-structure in the Fulani population has been already described [18,60] and has been linked to the Fulanisation phenomenon, i.e. the absorption of the sedentary local people within the nomadic Fulani groups and the consequent ethnicity shift of the former [63]. According to our data, FulaniB show a higher degree of admixture with neighboring populations, being more scattered in the PCA (Fig. 1 and Additional File 2) and showing different proportions of the ancestry components in the admixture and qpAdm analyses (Additional File 3, Supplementary figure S1 and S2, Additional File 1, Supplementary Table S4), as also highlighted by the D statistics (Additional File 1, Supplementary Table S3 and Additional File 3, Supplementary figure S3). When analyzing the ROH and IBD segments, the differences between the two Fulani clusters were confirmed, with FulaniA showing longer ROHs on average and a high degree of intra-population IBD sharing, probably a consequence of their endogamy. Interestingly, when dividing each cluster by region (i.e. Western Africa or Sahel), we could observe that the Sahelian FulaniA showed more and longer ROHs (Additional File 3, Supplementary figure S5), while Sahelian FulaniB were similar to other sub-Saharan people; on the other hand, FulaniA and FulaniB from western Africa were more similar (Additional file 3, Supplementary figure S5C). However, when considering short ROHs only, the pattern changed, replicating the FulaniA/FulaniB differentiation, as also observed from the IBD analysis (Additional file 3, Supplementary figure S5B and Additional file, Supplementary Table S5). All these observations seem to suggest that FulaniB experienced more admixture with other sub-Saharan groups in recent times, especially in the Sahel, possibly as a consequence of the Fulanisation. On the other hand, Sahelian FulaniA seems to have experienced stronger endogamy in recent times, leading to the presence of more and longer ROH segments, as also previously observed [15]. On the contrary, in western Africa, both clusters seem to show similar degrees of recent admixture. The reason for such internal population stratification is hard to explain since several aspects may have contributed to this (Fulanisation, social practices, urban vs. rural settlements [64]) and further studies on larger samples may be required to shed light on this.

## Conclusions

The population history of the Fulani, and the Saharan/Sahelian belt in general, is very complex and seems to have been strongly influenced by the major Holocene climate changes. In particular, the last Green Sahara (about 12000-5000 years ago) seems to have played a major role in the population dynamics of that period, allowing the contacts between different groups settled in that region, thus leading to the formation of pan-Saharan population(s). The exact ancestry composition of such ghost population(s) cannot be completely unveiled from modern genomes only; to the best of our knowledge, this is the first study consistently analyzing modern African whole genomes in the frame of ancient samples and this approach has been proved to be disproportionately informative, suggesting a similarity between putative ancient Saharans and Late Neolithic Moroccans. On this basis, these ancient Sahara group(s) may represent the ancestors of the modern Sahelian populations, including the Fulani, who may be the descendants of Saharan cattle herder groups who migrated westwards as a response to the aridification of the Sahara at the end of the Holocene climatic optimum. In the future, the analysis of more ancient specimens from western and central Sahel (if feasible) may shed new light on this hypothesized scenario, as well as on the western African ancestry of the Fulani. On the other hand, more whole genomes from modern African populations may represent an important resource to disclose different aspects of the history of the continent, for example to further understand the internal population structure of the Fulani people.

## Methods

### Sample selection and DNA quality control

We performed WGS on 43 individuals selected from our lab collection of DNA samples, obtained from peripheral blood, saliva or cultured cells. For each sample, geographic origin and ethnic affiliation were assigned solely on the basis of participants’ declarations and this information was used to select the individuals for WGS (Table 1 and Additional File 1, Supplementary Table S1).

WGS required specific quality and quantity parameters for the DNA to be analyzed: 1) quantity ≥ 3 μg; 2) absence or low amount of DNA degradation; 3) concentration ≥ 37.5 ng/μl; 4) purity, A260/280 = 1.8–2.0. First, more than 500 samples were initially screened to discard those below the quantity threshold. Then, degradation for the remaining samples was assessed by an electrophoretic run on a 1 % agarose gel. Concentration and purity of the samples passing the previous quality check were measured using a Qubit fluorometric quantification instrument (Thermo Fisher Scientific).

### WGS

Library preparation, sequencing, alignment and variant calling were performed by BGI-Tech (Hong Kong). WGS was performed on a DNBseq platform using the DNA Nanoballs (DNB) technology, after fragmentation of the genomic DNA, end repairs, ligation with adapters and ligation-mediated PCR of the 350 base pairs (bp) long fragments. Sequencing-derived image files were processed by DNBseq basecalling Software with default parameters to generate the sequence data of each individual as paired-end reads, stored in FASTQ format. Reads were aligned to the human reference genome (hg19/GRCh37) using the BWA software [65] and alignment files in BAM format will be produced using Picard [66]. The variant calling was performed using the HaplotypeCaller command of GATK [67], obtaining a list of raw variants in VCF format. The GATK Variant Quality Score Recalibration (VQSR), based on a machine learning algorithm, was used to discard spurious variants from the raw list. The final list of variants was listed in VCF format [68].

### Modern dataset

#### Selection of published whole genome sequences

In order to place our data in a broader context, we collected and analyzed 771 published WGS sequences, for a total of 814 individuals.

More specifically, our modern dataset includes samples from 5 different projects: 1) the samples from this study; 2) the samples from Fan et al. [19]; 3) a subset of relevant samples from the high coverage version of the 1000 Genomes Project [69]; 4) a subset of relevant samples from the Simons Genetic Diversity Project (SGDP) [30]; 5) a subset of relevant samples from the whole genomes from Human Genetic Diversity Project (HGDP) [31]. For all the datasets, the starting files were VCFs.

#### Liftover

Sequencing data from two published datasets were aligned against the hg38/GRCh38 human reference sequence [31,69]. In order to make the variant list comparable, these VCF data were converted to the hg19/GRCh37 coordinates using the LiftoverVcf tool in Picard [66].

#### Sample selection for SGDP, HGDP and 1000 Genomes project

The variant datasets from the three worldwide variation projects [30,31,69] are available as multi-sample VCFs, however only a subset of samples was relevant for our study. For the selection, we focused on different ethnic groups and geographical regions in Africa, Europe and Asia. Moreover, since some populations were over-represented with a large sample size, we randomly selected 25 individuals from each large population in each dataset to make their sample size comparable with the number of Fulani individuals (25). After these selection steps, the samples to be excluded were discarded using bcftools [70]. These steps have been performed on each dataset separately.

#### African datasets

The two African datasets, i.e. the one analyzed in this study and the one from Fan et al. [19], were organized with one VCF for each sample. First, we had to create a multisample VCF for each dataset, merging the samples with bcftools. Since each individual may have private variants, this can create a large amount of missing data when merging different individuals. However, considering the high quality of the sequencing data, we can reliably state that the positions not listed in the individual VCFs show the reference state in the respective samples and so they have been coded as homozygous reference during the merging step using the-0 option in the bcftools view command [70].

#### Data merging and SNP filtering

The 5 datasets have been first analyzed in two batches: 1) datasets originally in the hg19/GRCh37 assembly, i.e. this study, SGDP and Fan et al. [19,30]; 2) datasets originally in hg38/GRCh38 and then converted by liftover, i.e. HGDP and 1000 Genomes project [31,69].

The datasets in each batch have been merged with bcftools [70], setting variants that were not present in one or more datasets as missing in the corresponding individuals. Then, only the biallelic SNPs on the autosomes have been kept and the strict callable mask from the 1000 Genome Project [29] was applied. After this step, the two batches have been merged together using bcftools [70], averaging the depth (DP) and mapping quality (MQ) values across all the samples to create a single multisample VCF. In order to discard false variant calls, we applied the following filters: 1) minor allele frequency ≥ 1%; 2) QUAL > 30; 3) average MQ > 40, leaving more than 12 million variant calls.

#### Final modern datasets

The modern dataset obtained in the previous steps was then further filtered to create two different datasets used for the haplotype-based or frequency-based analyses.

For the haplotype-based methods, the modern dataset was filtered to discard variants missing in at least one sample using the --geno 0 option in plink [71], for a total of 4143934 SNPs.

For the allele frequency based analyses, the modern dataset has been filtered using plink [71] to discard the variants in linkage disequilibrium (LD) and missing in more than 10% of samples. The LD filter was applied using the --indep-pairwise command with a window size of 200, shifting by 25 SNPs and a maximum pairwise r^2^ threshold of 0.5, while the missingness filter was applied using the --geno 0.1 option, for a total of 1907678 SNPs.

#### Ancient and modern dataset

We merged our LD-pruned modern dataset with the modern and ancient genome-wide SNP data available from the Allen Ancient DNA Resource (AADR, V44.3) [40] using plink [71]. First, we discarded the AADR samples that were present also in our modern dataset from the published genetic projects [19,30,31,69] and kept only individuals from the same geographic areas as the modern dataset (Additional File 2, Supplementary Table S2). Then, the two sets of data were merged based on common variants, for a total of 226222 SNPs in 2615 samples. Additional ancient samples to be used as comparison or outgroups have been added for specific analyses, as explained in the corresponding method sections.

#### PCA

Principal component analysis (PCA) was performed on our LD-pruned modern dataset using the smartpca tool from EIGENSOFT version 7.2.0 [72]. We also performed the PCA on the modern + ancient dataset by estimating the principal components from modern individuals only and projecting the ancient samples onto such components.

#### Admixture

Patterns of population structure were explored performing a model-based individual ancestry analysis using ADMXITURE v1.3.0 [44], running 10 independent iterations for K ancestral clusters ranging from 2 to 10 with the unsupervised approach for the modern dataset. For the modern + ancient dataset, the modern samples were first pseudohaploidized to make them consistent with the ancient pseudohaploid data. Then, the ADMIXTURE analysis was performed as described above.

#### f statistics, qpWave/qpAdm and admixture graph

The D statistics have been estimated using ADMIXTOOLS version 7.0.1 [73] on the modern + ancient dataset after adding the chimpanzee and Neanderthal data (for a total of 2626 individuals) from the AADR (V44.3) [40] to be used as outgroups. In order to describe the ancestry of Fulani samples as a combination of prehistoric/modern groups, we used the qpWave/qpAdm framework. In detail, for each Fulani group, given a set of left and right populations, and for the number of left sources K {2,3}: a) we evaluated if the right samples can be used to significantly discriminate the provided sources, using qpWave. Right populations present in sources were excluded for that specific test; b) if step a) provided a significant *p* value < 0.01, we used qpWave to evaluate if the target population could be described as a combination of K sources; c) if b) provided a *p* value > 0.01, we used qpAdm to describe the target as a mixture of the K sources employed for that specific experiment. Given the high rate of missingness in some of our tested samples we used the option ‘‘allsnps=YES’’. As left populations, we used: “GWD”, “MSL”, “Cameroon_SMA_6000-3000BCE”, “Bantu”, “Mada”, “Bulala”, “Agew”, “ESN”, “Berbers”, “Morocco_EN_8000-6000BCE”, “Morocco_LN_6000-3000BCE”, “Bedouin”, “Turkey_N_8000-6000BCE”. As right populations, we used: “San”, “Czech_Vestonice”, “Spain_ElMiron_before10000BCE”, “Italy_North_Villabruna_HG_before10000BCE”, “WHG”, “ONG.SG”, “Han”, “Papuan.SDG”, “Russia_Ust_Ishim_HG_published.DG”, “Russia_Kostenki”, “Russia_MA1_HG.SG”, “Chimp.REF”, “Neanderthal”, “Karitiana.SDG”, “Iran_GanjDareh_N_8000-6000BCE”.

For the admixture graph, we harnessed the function “graph_Auto”, implemented in admixtools 2 [74], to model the evolutionary history and admixture edges of each Fulani group, together with a set of representative populations (“Morocco_Iberomaurusian_before10000BCE”, “Mbuti”,“MSL”,,“Morocco_LN_6000-3000BCE”, “Yoruba”, “Altai_Neanderthal.DG”, “Chimp.REF”, “Israel_Natufian_before10000BCE”, “Iran_GanjDareh_N_10000-8000BCE”). We estimated graphs varying the number of admixture edges from 2 to 5, and repeated the whole process 10 times. For each admixture edge the three most supported graphs were retained and plotted.

#### Identity-by-descent and runs of homozygosity

The identity-by-descent (IBD) analysis has been performed on the non-LD-pruned modern dataset using the software IBIS version 1.20.6 [75] with the default parameters. The runs of homozygosity (ROH) analysis was performed on the same dataset using plink after slightly modifying the parameters previously used for Sahelian populations [59] as follows: --homozyg-snp 30 --homozyg-kb 300 --homozyg-density 30 --homozyg-window-snp 30 --homozyg-gap 1000 --homozyg-window-het 1--homozyg-window-missing 15 --homozyg-window-threshold 0.05. The raw outputs have been manipulated and plotted using the ggplot2 and dplyr packages in R version 4.1.3.

### Additional files

**Additional file 1: Table S1:** Description of the samples in the modern dataset. The language information has been reported only for African populations (AA: Afro-Asiatic, K: Khoisan, NC: Niger-Congo, NS: Nilo-Saharan). **Table S2:** Description of the additional (modern and ancient) samples included in the modern + ancient dataset from Allen Ancient DNA Resource (AADR, V44.3). **Table S3:** D-statistics in the form D(Fulani,sub-Saharan Africa,Eurasia/North Africa,Chimpanzee). **Table S4:** qpAdm models of Fulani groups with two or three putative sources. **Table S5:** Identity-by-descent (IBD) segments by Population group (see Table S1). The Fulani clusters have not been divided by Region (Sahel or WAfrica). Genetic length is reported in cM. Short IBDs: segments shorter than 5 cM; Long IBDs: segments longer than 5 cM. norm: normalization by the number of possible comparisons. **Table S6:** D-statistics in the form D(Sahel, West Africa, Fulani, Chimpanzee). **Table S6:** Runs of homozygosity (ROH) by Population group (see Table S1). ROH length is in kb. Short ROHs < 1.5 Mb, Long ROHs > 1.5 Mb. (XLSX)

**Additional file 2:** Interactive version of the PCA in figure 1 based on the modern dataset. (HTML)

**Additional file 3: Supplementary Figure S1:** Full plot of the admixture analysis of the modern dataset from K=2 to K=10. The classification of samples by region is reported at the bottom, the full population list is reported in the table on the right in the same order as in the plot (from left to right). **Supplementary Figure S2:** Full plot of the admixture analysis of the modern + ancient dataset from K=2 to K=10. The classification of samples by region is reported at the bottom, the full population list is reported in the table on the right in the same order as in the plot (from left to right). **Supplementary Figure S3:** Plot of the D statistic in the form D(Fulani, Mandenka, Eurasia/Northern Africa, Chimpanzee), where the two clusters FulaniA and FulaniB are represented separately (blue and purple respectively). Only the significant values (|Z| > 3) are plotted. **Supplementary Figure S4:** Admixture graphs with edges from 2 to 5, from top to bottom. For each tested edge, the best three graphs with the lowest score have been retained and plotted. **Supplementary Figure S5:** Scatterplot of the runs of homozygosity (ROH) segments by population comparing the mean number of ROH with their mean length. A: all ROH segments; B: short ROH segments (< 1.5 Mb); C: long ROH segments (> 1.5 Mb). (PDF)

**Additional file 4:** Interactive version of the PCA based on the modern + ancient dataset. (HTML)

### Abbreviations

AA: Afro-Asiatic language
AADR: Allen Ancient DNA Resource
AN: Anatolian Neolithic
BCE: Before Common Era
BP: Before Present
CHG: Caucasus Hunter-Gatherers
DNB: DNA Nanoballs
DP: depth
HGDP: Human Genome Diversity Project
IBD: Identity-by-descent
LD: linkage disequilibrium
MQ: mapping quality
mtDNA: mitochondrial DNA
NC: Niger-Congo language
NS: Nilo-Saharan language
PC: Principal component
PCA: Principal component analysis
ROH: Runs of homozygosity
SGDP: Simons Genome Diversity Project
SNP: single nucleotide polymorphism
WGS: Whole genome sequencing
WHG: Western Hunter-Gatherers

## Declarations

### Ethics approval and consent to participate

All the DNA samples analyzed here have been selected from our lab collection, collected on a voluntary basis over the past decades after obtaining an appropriate informed consent and according to all the regulations in effect at that time. The use of this “historical” collection in genomic studies was approved by the Ethical Committee Fondazione IRCCS Policlinico San Matteo (protocol number 0028298/22).

### Availability of data and materials

The VCF files of the samples generated during this study are available through the European Genome-phenome Archive (EGA, https://ega-archive.org/) under the accession number [.] (files available upon publication).

## Funding

This work was supported by National Geographic (Grant HJ-139R-17 to FC) and by Sapienza University of Rome (grant RG120172B7F263BA to FC). BB was supported by a postgraduate scholarship from the Istituto Pasteur—Fondazione Cenci Bolognetti.

### Authors’ contributions

FC and ED conceived the study. FC, BT, DAS, GC, OS, GDB and PA collected and provided the DNA samples. ED, BB and SO selected the DNA samples to be analyzed in this study. ED assembled the modern dataset from new and published sequencing data. FRi and FRa assembled the modern + ancient dataset. ED, FRi, FRa, FM, MH, LP, MB and BT performed the bioinformatic analysis. ED, FC, BT, FM, MM and KT discussed and interpreted the results. ED wrote the original draft with inputs from the other authors. All the authors revised and approved the final version of the manuscript.

## Supporting information

Modern PCA interactive

Ancient-modern PCA interactive

## Acknowledgements

The authors are grateful to all the donors for providing DNA samples and to the people that contributed to the sample collection. In particular, they thank Prof. Gabriella Spedini for her help in the collection of the samples. Analyses were carried out using the facilities of the High Performance Computing Center of the University of Tartu.

