## Supplementary Figures for "Echoes from the last Green Sahara: whole genome analysis of Fulani, a key population to unveil the genetic evolutionary history of Africa"

**Supplementary Figure S1**: Full plot of the admixture analysis of the modern dataset from K=2 to K=10. The classification of samples by region is reported at the bottom, the full population list is reported in the table on the right in the same order as in the plot (from left to right).

**Supplementary Figure S2**: Full plot of the admixture analysis of the modern + ancient dataset from K=2 to K=10. The classification of samples by region is reported at the bottom, the full population list is reported in the table on the right in the same order as in the plot (from left to right).

**Supplementary Figure S3**: Plot of the D statistic in the form D(Fulani, Mandenka, Eurasia/Northern Africa, Chimpanzee), where the two clusters FulaniA and FulaniB are represented separately (blue and purple respectively). Only the significant values ($|Z| > 3$) are plotted.

**Supplementary Figure S4**: Admixture graphs inference of Fulani together with a set of representative populations. For each admixture edge from 2 to 5, the best three graphs with the lowest score have been retained and plotted, from top to bottom.

**Supplementary Figure S5**: Scatterplot of the runs of homozygosity (ROH) segments by population comparing the mean number of ROH with their mean length. A: all ROH segments; B: short ROH segments (< 1.5 Mb); C: long ROH segments (> 1.5 Mb).

### Supplementary Figure S1

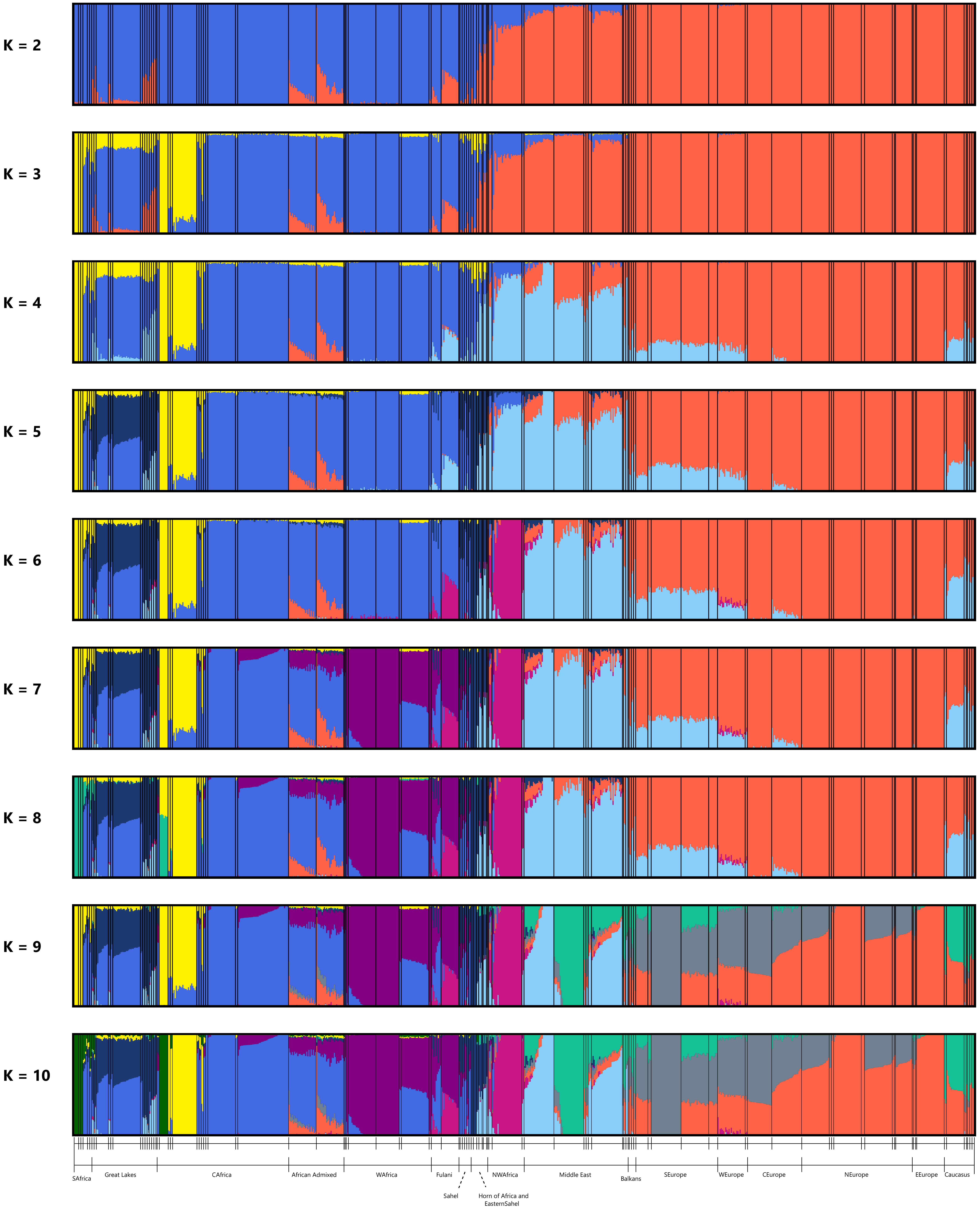


| Regions | Populations |
|---|---|
| SAfrica | Jhn |
| | Khm |
| | San |
| | Bnt |
| | Her |
| | Tsw |
| Great Lakes | Snd |
| | Hdz |
| | Bnt |
| | Kky |
| | Lwk |
| | LWK |
| | Luo |
| | Msi |
| | Ogk |
| | Sgw |
| | Elm |
| | Irw |
| | Rnd |
| | Som |
| CAfrica | Kba |
| | Mbt |
| | Bka |
| | Bkl |
| | Bik |
| | Lmd |
| | Ngb |
| | Bdz |
| | Tks |
| | Esn |
| | ESN |
| | Igb |
| | Yrb |
| African Admixec | ACB |
| | ASW |
| WAfrica | Akn |
| | Wli |
| | Gwd |
| | GWD |
| | Mnk |
| | Msl |
| | MSL |
| | Wlf |
| Fulani | Fulani B |
| | Fulani A |
| Sahel | Fal |
| | Mda |
| | Mss |
| | Sng |
| | Bla |
| | Lka |
| Horn of Africa and East. Sahel | Mur |
| | Dnk |
| | Aar |
| | Amh |
| | Tgy |
| | Agw |
| | Sho |
| NWAfrica | Ber |
| | Mor |
| | Mzb |
| | Shw |
| Middle East | Bed |
| | Dru |
| | Irn |
| | Iqj |
| | Jrd |
| | Pal |
| | Smr |
| | Trk |
| | Ymj |
| Balkans | Alb |
| | Bul |
| | Cre |
| | Gre |
| SEurope | Brg |
| | Ita |
| | Sar |
| | TSI |
| | Tsi |
| WEurope | IBS |
| | Ibs |
| CEurope | Bsq |
| | Fre |
| NEurope | CEU |
| | Eng |
| | Est |
| | FIN |
| | Fin |
| | GBR |
| | Ice |
| | Nrw |
| | Orc |
| EEurope | Cze |
| | Hun |
| | Pol |
| | Rus |
| Caucasus | Abk |
| | Ady |
| | Arm |
| | Chc |
| | Geo |
| | Lzg |
| | Nos |

### Supplementary Figure S2

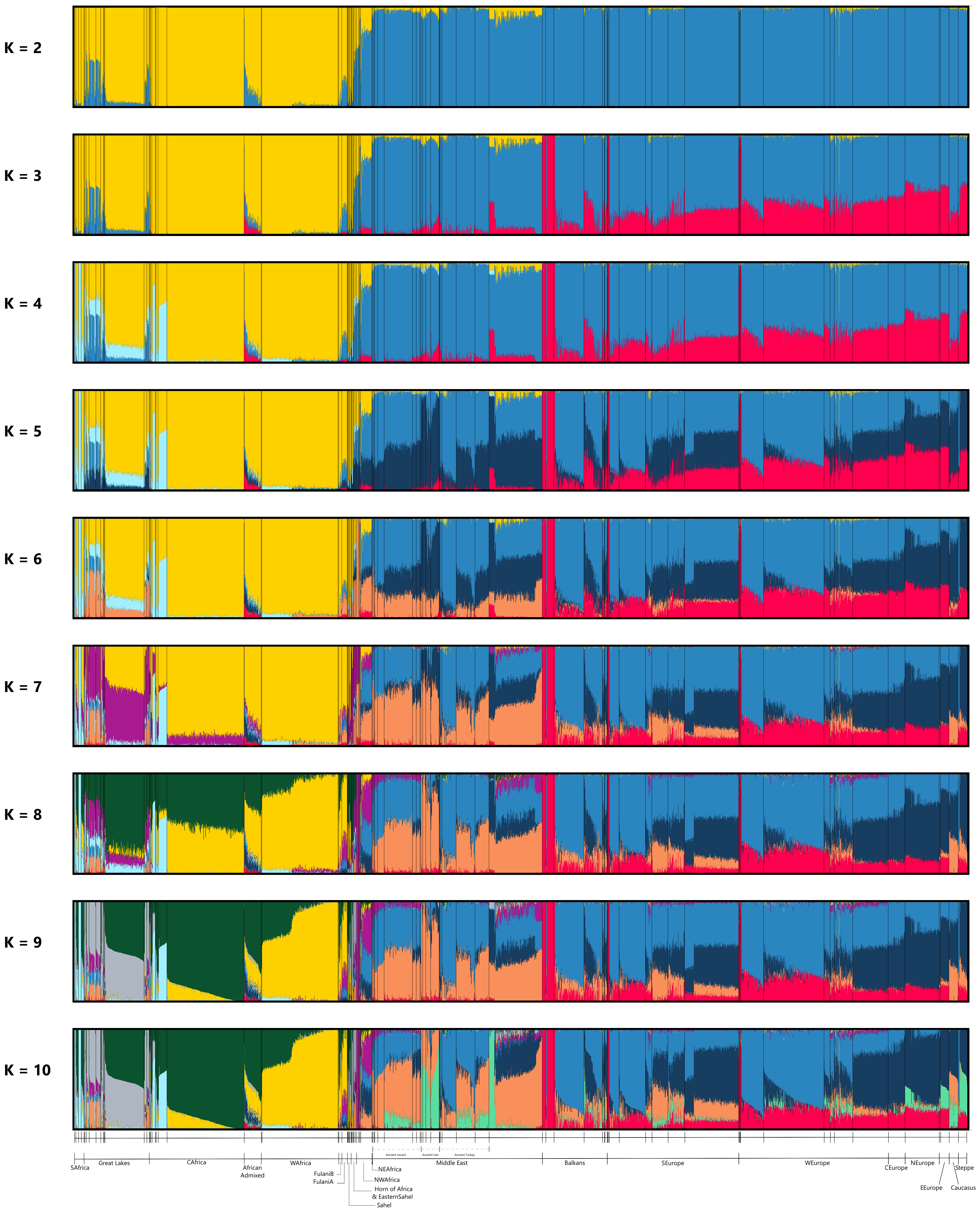


| Region | Population |
|---|---|
| SAfrica | 300BC.500AD |
| | 500.1900AD |
| | Khoisan |
| | NC.Bantu |
| Great Lakes | 8000.6000BC |
| | 6000.3000BC |
| | 3000.1000BC |
| | 1000.300BC |
| | 300BC.500AD |
| | 500.1900AD |
| | Khoisan |
| | Hadza |
| | NC.Bantu |
| | NS.Nilotic |
| | AA.Cushitic |
| CAfrica | 6000.3000BC |
| | 3000.1000BC |
| | 500.1900AD |
| | NS.Sudanic |
| | NC.Adamawa |
| | NC.Bantoid |
| | NC.Bantu |
| | NC.BenueCongo |
| African Admixed | African Admixed |
| WAfrica | NC.VoltaCongo |
| | NC.Mende |
| | NC.Wolof |
| Fulani | Fulani B |
| | Fulani A |
| Sahel | NC.Adamawa |
| | NC.Mada |
| | AA.Chadic |
| | NS.Songhai |
| | NS.Sudanic |
| Horn of Africa&Eastern Sahel | 3000.1000BC |
| | NS.Sudanic |
| | NS.Nilotic |
| | AA.Omotic |
| | AA.Semitic |
| | AA.Cushitic |
| NWAfrica | before10000BC |
| | 8000.6000BC |
| | 6000.3000BC |
| | AA.Berbers |
| NEAfrica | 1000.300BC |
| Middle East | Levant_before10000BC |
| | Levant_10000.8000BC |
| | Levant_8000.6000BC |
| | Levant_6000.3000BC |
| | Levant_3000.1000BC |
| | Levant_1000.300BC |
| | Levant_300BC.500AD |
| | Levant_500.1900AD |
| | Iran_10000.8000BC |
| | Iran_8000.6000BC |
| | Iran_6000.3000BC |
| | Iran_3000.1000BC |
| | Iran_1000.300BC |
| | Turkey_before10000BC |
| | Turkey_10000.8000BC |
| | Turkey_8000.6000BC |
| | Turkey_6000.3000BC |
| | Turkey_3000.1000BC |
| | Various Modern |
| Balkans | 10000.8000BC |
| | 8000.6000BC |
| | 6000.3000BC |
| | 3000.1000BC |
| | 300BC.500AD |
| | 500.1900AD |
| | Various Modern |
| SEurope | before20000BC |
| | before10000BC |
| | 10000.8000BC |
| | 6000.3000BC |
| | 3000.1000BC |
| | 1000.300BC |
| | 300BC.500AD |
| | 500.1900AD |
| | Various Modern |
| WEurope | before10000BC |
| | 10000.8000BC |
| | 8000.6000BC |
| | 6000.3000BC |
| | 3000.1000BC |
| | 1000.300BC |
| | 300BC.500AD |
| | 500.1900AD |
| | Various Modern |
| CEurope | CEurope |
| NEurope | NEurope |
| EEurope | EEurope |
| | Russian |
| Caucasus | Various Modern |
| Steppe | Yamnaya |

### Supplementary Figure S3

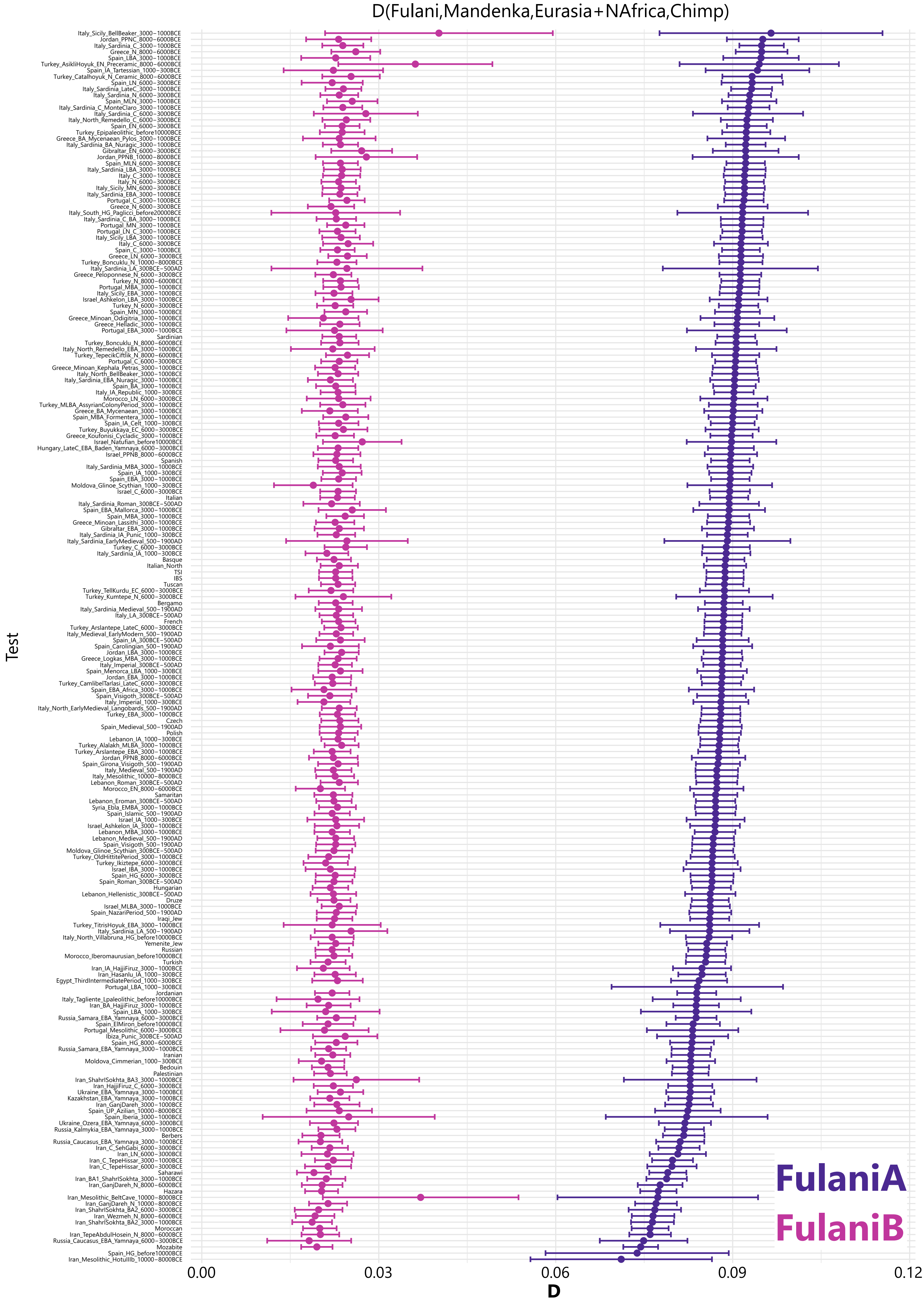

#### Supplementary Figure S4

Edge = 2

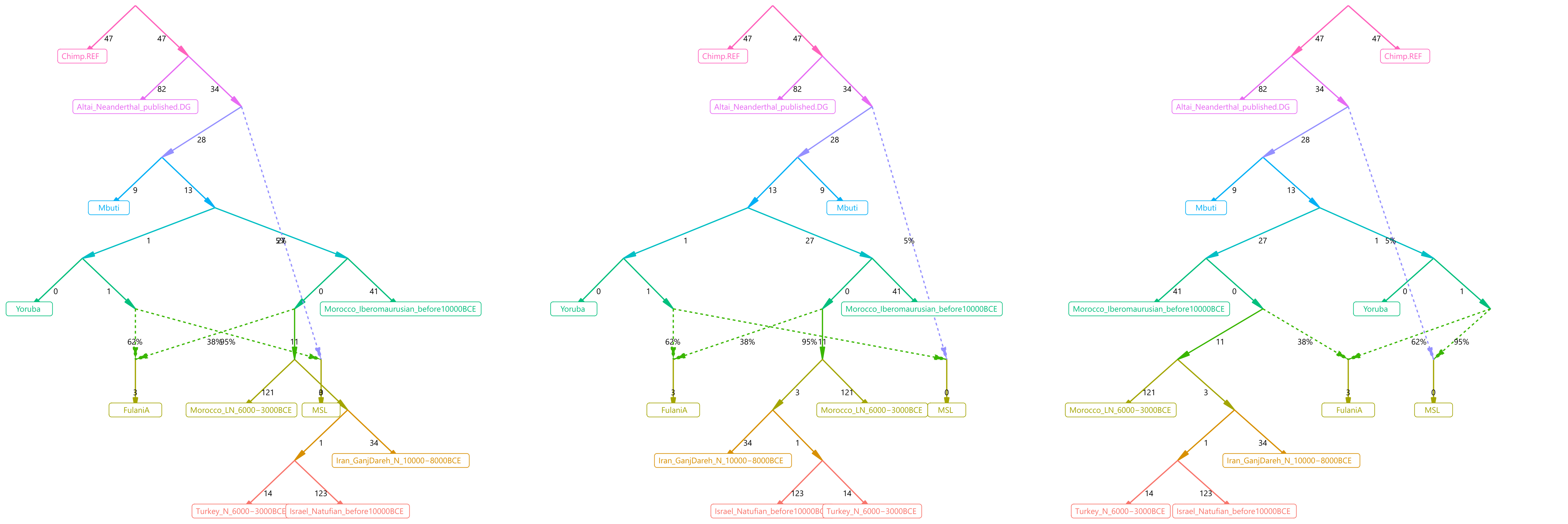


Edge = 3

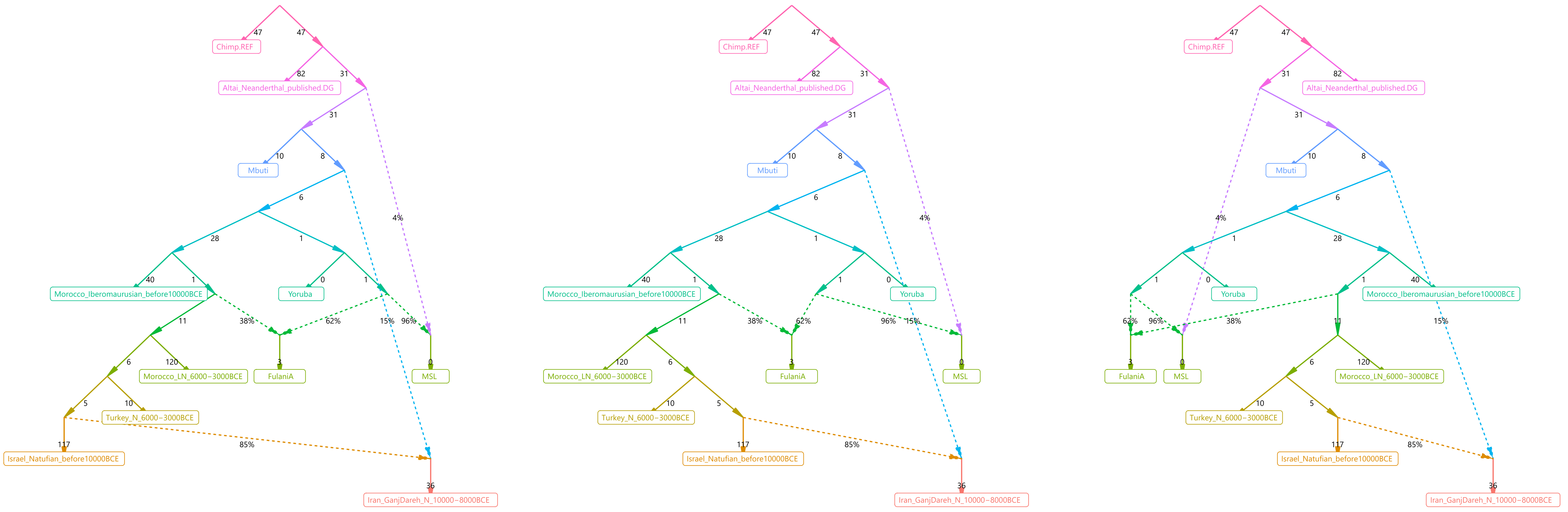


Edge = 4

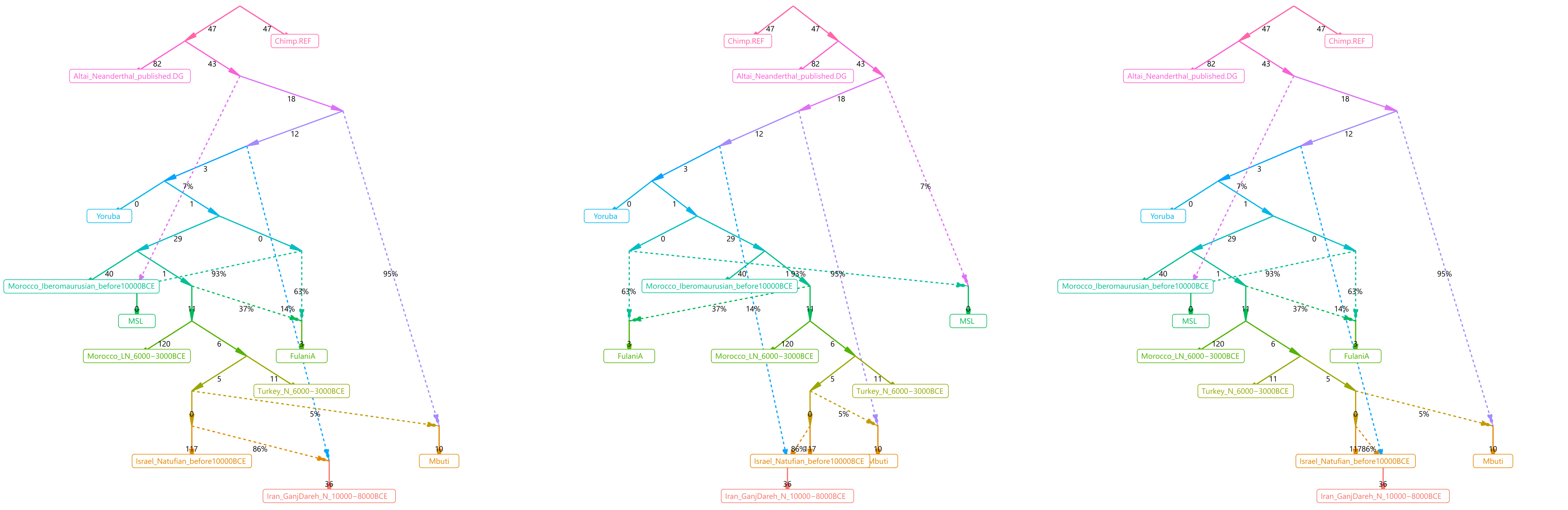


Edge = 5

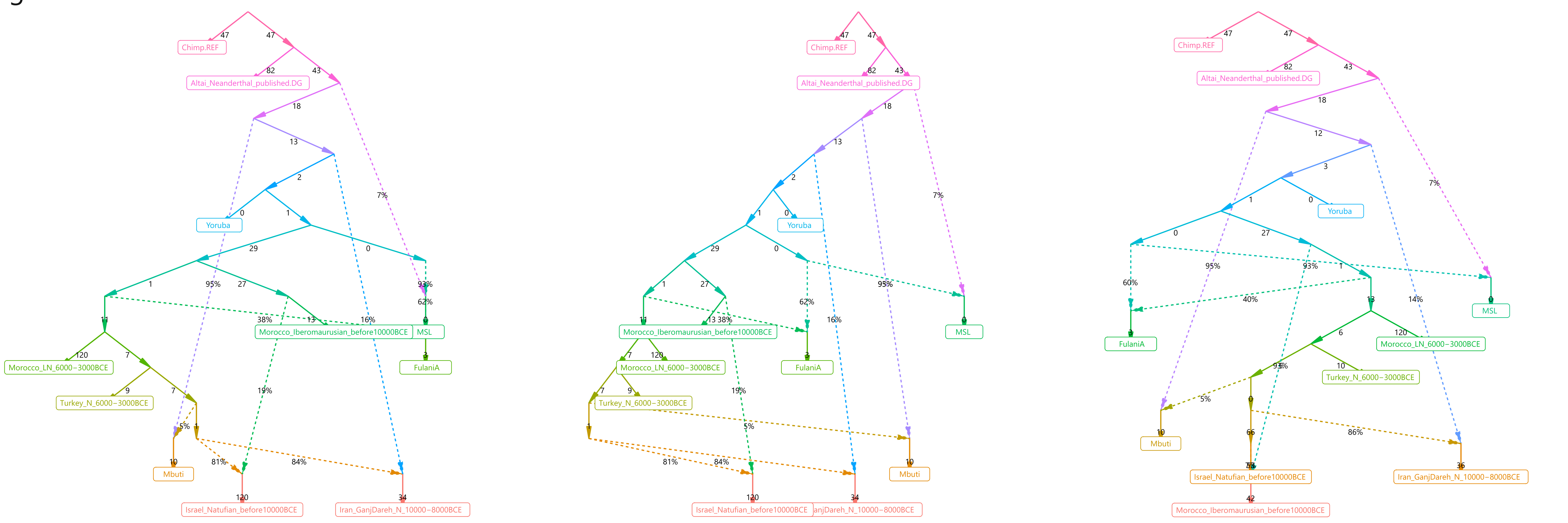

### Supplementary Figure S5

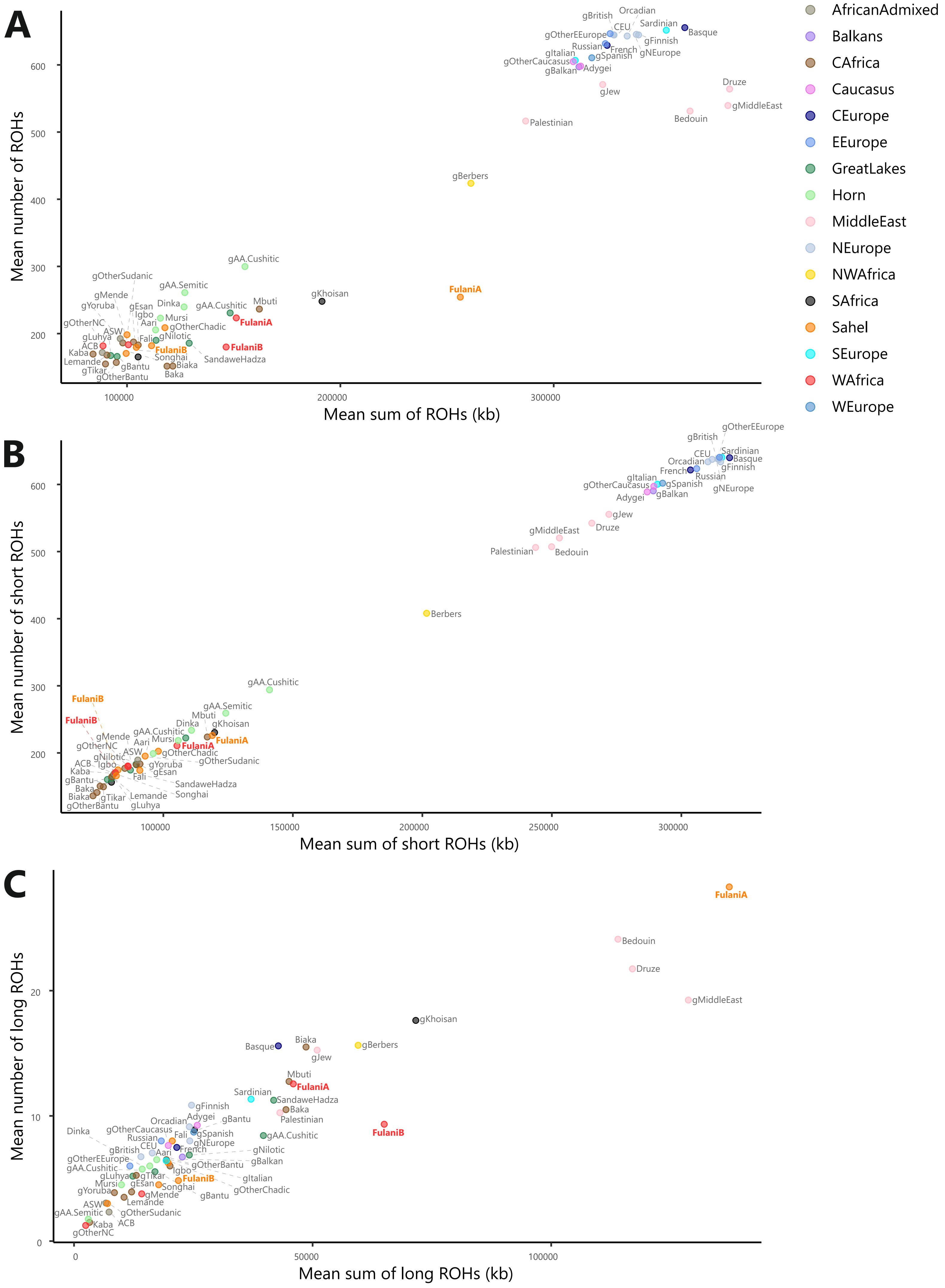
